# Dietary zinc restriction mimics protein restriction and extends lifespan in *Drosophila*

**DOI:** 10.1101/2024.08.28.610190

**Authors:** Hina Kosakamoto, Hide Aikawa, Souto Kitazawa, Chisako Sakuma, Rina Okada, Masayuki Miura, Fumiaki Obata

## Abstract

Dietary restriction extends lifespan in model organisms, mainly through dietary amino acids. Compared to macronutrients, the effect of dietary micronutrients on organismal lifespan has not been intensively investigated. Here, using a synthetic diet, we test whether restriction of each micronutrient, including vitamins and minerals, affects lifespan and fecundity in adult *Drosophila*. While restriction of many of these micronutrients have either negative or no impact on lifespan, zinc (Zn) restriction alone can increase it. Dietary Zn restriction (ZnR) decreases fecundity, increases starvation resistance, and promotes preference for feeding amino acids, in adult females, phenocopying dietary amino acid restriction. Our study demonstrates that dietary intake of trace elements has profound impacts on physiology and lifespan, and that limiting dietary zinc may be a strategy to improve the healthspan of animals.

## Introduction

Nutrition is actively involved in homeostasis and determines healthspan of animals. Dietary restriction is known to increase lifespan in various organisms and its mechanism has been intensively studied^1–3^. How dietary factors involves lifespan extension has been analysed and essential amino acids are shown to be the key nutrients determining lifespan^4,5^. However, compared to macronutrients, how dietary micronutrients, such as minerals and vitamins, influence lifespan and healthspan is not studied well.

A synthetic diet, or holidic medium, is readily available in *Drosophila* and is useful for manipulating individual nutrients^5,6^. It is used to perform *loss-of-function* experiments to elucidate the effect of a single nutrient on metabolism, tissue homeostasis, behaviour, fecundity, and lifespan. The synthetic diet contains approximately forty pure ingredients, including a sugar, lipids, amino acids, vitamins, and minerals. Among the micronutrients, some trace elements, as well as vitamins, are nutritional requirements for most animals. The synthetic diet contains four trace (heavy) metals: zinc (Zn), copper (Cu), iron (Fe), and manganese (Mn). Depletion of all metal ions can slightly increase lifespan and decrease fecundity^5^. To date, the effect of restriction of each metal on metabolism, fecundity, and lifespan have not been examined well.

In this study, we addressed how dietary restriction of each micronutrient such as vitamins or metals can influence *Drosophila* lifespan by utilising the synthetic diet. We found that restriction of Zn, but not Cu, Fe, Mn or vitamins, can extend lifespan. Zinc restriction (ZnR) decreases fecundity, increases starvation resistance, and promotes preference to amino acid feeding, suggesting that it phenocopies typical dietary amino acid restriction.

## Results

### Dietary Zinc restriction extends lifespan

For dietary manipulation of micronutrients, we prepared synthetic diets (please see Methods for detail). To test the effect of trace metals and vitamins, we fed female Canton-S flies with the diet which lacks each one of vitamins or heavy metals (Fig. 1a). Of note, the flies were raised using a standard yeast-based diet, which contained all these micronutrients, to avoid developmental lethality, then flip the adult female flies to the synthetic diets at day 2 post eclosion (Fig. 1a, Table 1). We assumed that restriction of an essential nutrient throughout adult life would lead to early mortality. As we expected, dietary restriction of many vitamins shortened organismal lifespan (Fig. 1b). In contrast, complete drop out of each metal ion did not strongly shorten lifespan, while dietary restriction of Fe slightly increased lifespan (Fig. 1c). Notably, dietary Zn restriction (ZnR) significantly extended lifespan (Fig. 1c). Since it was intriguing to see the specific and striking effect of Zn on lifespan, we decided to focus on this micronutrient for the rest of the study.

**Fig. 1.**
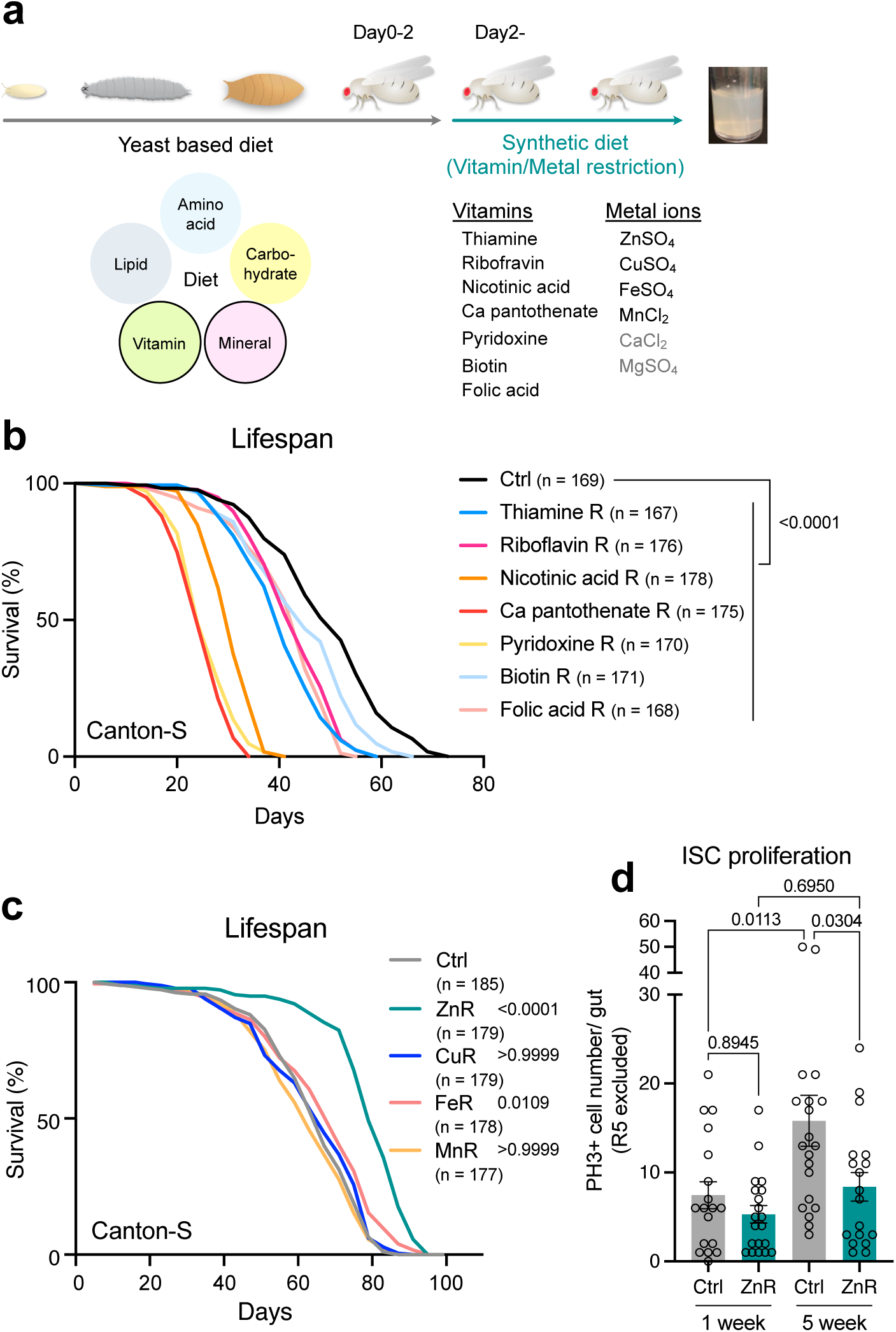
Zinc restriction extends female lifespan and gut healthspan. **a,** The scheme of dietary manipulation. Each of seven vitamins and four trace metals was dietary restricted from day 2. **b,c,** Lifespans of female Canton-S flies fed with or without vitamin- (**b**) or metal- (**c**) restricted diets. Sample sizes (n) are shown in the figure. For the statistics, a log-rank test was used. **d,** Intestinal stem cell proliferation indicated by phospho-histone H3 positive cells in the guts of female Canton-S flies. The flies were fed the synthetic diet with or without zinc for one or five weeks. n = 18 (1-week Ctrl and 5-week ZnR) or 20 (1-week ZnR and 5-week Ctrl). For the statistics, one-way ANOVA with Holm-Šídák’s multiple comparison test was used. For the graph, the mean and SEM are shown. Data points indicate biological replicates. Source data are provided as a Source Data file.

**Table 1.**
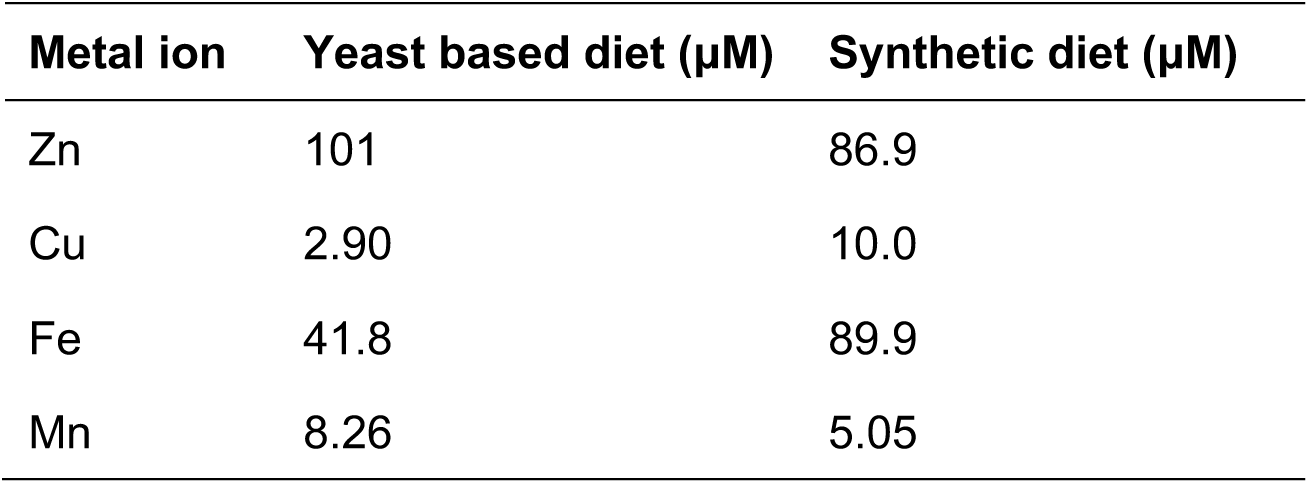
Metal ion content in fly diets.

| **Metal ion** | **Yeast based diet ($\mu\text{M}$)** | **Synthetic diet ($\mu\text{M}$)** |
| --- | --- | --- |
| Zn | 101 | 86.9 |
| Cu | 2.90 | 10.0 |
| Fe | 41.8 | 89.9 |
| Mn | 8.26 | 5.05 |

Substantial evidence indicated that ISC hyperproliferation is a hallmark of *Drosophila* ageing, at least of the gut, that number of which inversely correlates with lifespan and healthspan^7–10^. We found that ZnR suppressed the increased ISC division during ageing, suggesting that gut homeostasis is maintained by ZnR (Fig. 1d). Together, these data suggested that ZnR extends organismal lifespan and increases intestinal healthspan.

### Dietary Zinc restriction decreases internal Zinc levels

The profound positive impact of Zn, but not other metals, restriction on female lifespan implied that, in physiological conditions, internal Zn levels might be specifically elevated with age, promoting pathology of organismal ageing. We measured the level of four metals in the whole body of young (day5) and aged (day42) flies reared on the standard yeast-based diet, by using inductively coupled plasma-mass spectrometry (ICP-MS). The analysis measured both of free and protein-bound metal ions. The level of Zn was indeed increased in the aged flies (1.7-fold increase) (Fig. 2a). However, this tendency was also the case for other metals, suggesting specificity of lifespan extension by ZnR is not due to the specific accumulation of internal metal levels (Fig. 2b-d). We then quantified how strongly Zn levels are decreased in female flies fed with ZnR diet. The average Zn levels in the whole body of female flies after one-week of ZnR was 13.8 ng/mg body weight, compared to the control 112 ng/mg body weight, indicating that whole body Zn levels decreased to 12% of the control within a week (Fig. 2e).

**Fig. 2.**
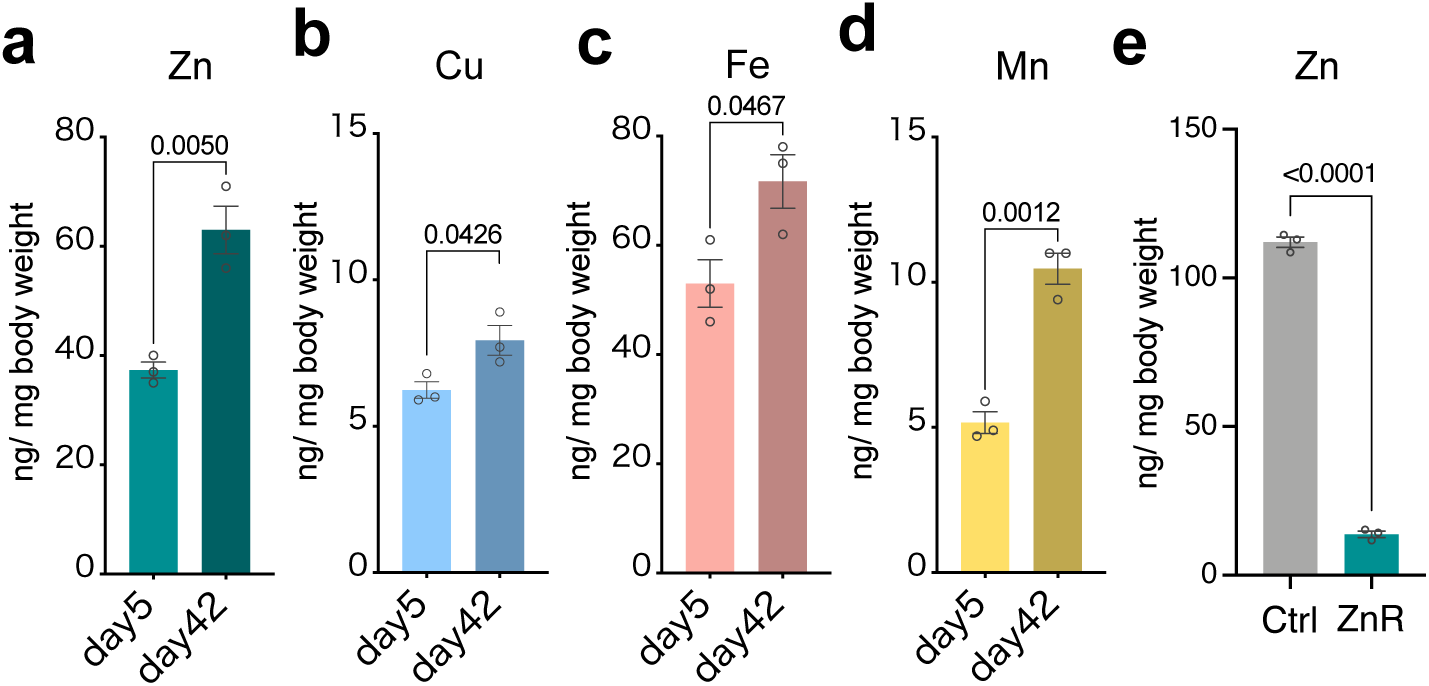
Internal levels of metal ions during ageing and dietary restriction. **a-d,** Internal levels of zinc (**a**), copper (**b**), iron (**c**), and manganese (**d**) in female Canton-S flies at day 5 and 42 reared on a standard yeast-based diet. n = 3. For the statistics, a two-tailed Student’s *t* test was used. **e,** Internal levels of zinc in female Canton-S flies fed the synthetic diet with or without zinc. n = 3. For the statistics, a two-tailed Student’s *t* test was used.

It is worth noting that the Zn concentrations of the flies fed with the complete synthetic diet were higher than those in the standard yeast-based diet (Fig. 2a,e). However, these diets contained approximately the same amount of total Zn levels (Table 1). This discrepancy is probably due to the bioavailability as Zn in the standard diet is likely chelated, while that in the synthetic diet is free ions. Difference in the Zn levels in these two dietary conditions would impact the physiology and lifespan of flies, however it is very difficult to compare them as synthetic diet contains minimal number of nutrients. Empirically, however, higher Zn levels in the flies fed with the control synthetic diet do not show the shorter lifespan compared to that with standard yeast diet.

### Small amount of dietary Zinc is necessary for lifespan extension

Given that complete Zn restriction does not enhance mortality, we suspected that Zn was contaminated in some components of the synthetic diet and provided minimal Zn requirement for flies to maintain homeostasis. Especially, an agar can contain various trace elements. To check this, we quantified heavy metals in agar or agarose obtained from several manufacturers. Indeed, the agar we used for the synthetic diet has relatively higher amount of Zn as well as other trace elements (1,523 ng/g for Zn, 164 ng/g for Cu, 68,994 ng/g for Fe, 8,261 ng/g for Mn) (Fig. 3a-d). Among tested, we found that an agarose from Nippon Gene Inc. has minimum amount of contaminated Zn (15 ng/g for Zn, 350 ng/g for Cu, 808 ng/g for Fe, 46 ng/g for Mn) (Fig. 3a-d). Therefore, we analysed lifespan of female flies under synthetic diets made with this agarose. ZnR using the agarose failed to increase lifespan and rather slightly decrease it (Fig. 3e), suggesting that the lifespan extension by ZnR demands a small amount of contaminated Zn in the agar and that harsher ZnR has thus negative impact on longevity.

**Fig. 3.**
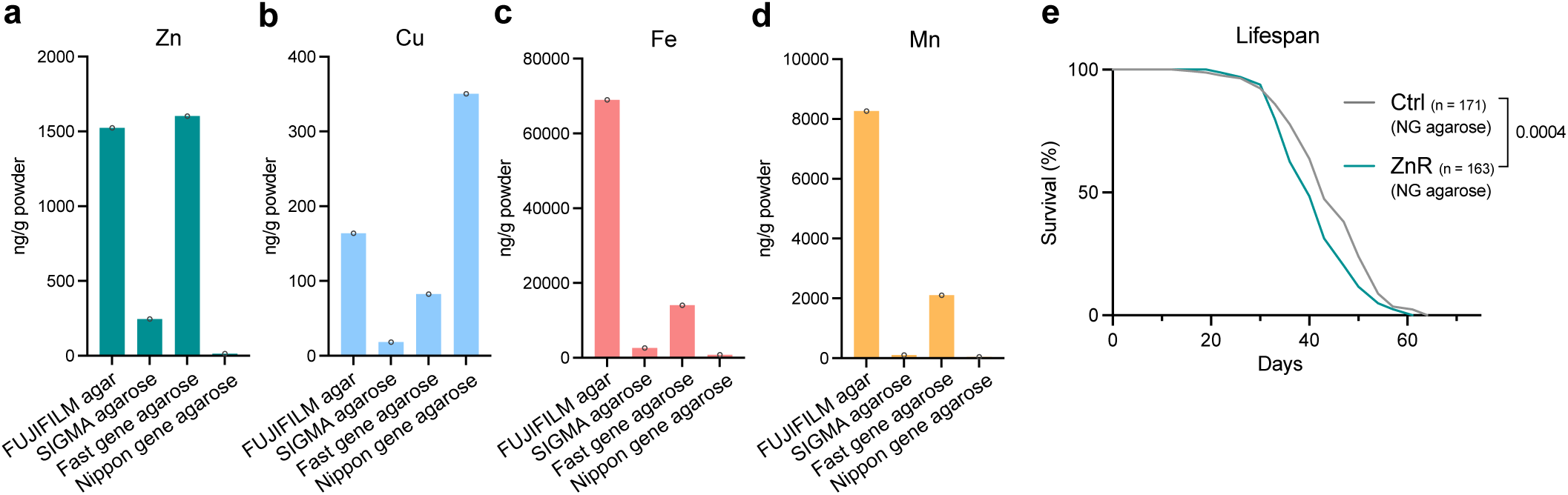
Zinc restriction with a minimum zinc contamination shortens lifespan. **a-d,** The content of zinc (**a**), copper (**b**), iron (**c**), and manganese (**d**) in the agar or agarose powder. n = 1. **e,** Lifespans of female Canton-S flies fed with or without a zinc-restricted diet using Nippon gene agarose instead of FUJIFILM agar. Sample sizes (n) are shown in the figure. For the statistics, a log-rank test was used.

### Zinc restriction phenocopies dietary yeast restriction

In general, dietary restriction induces the trade-off between fecundity and lifespan. As we expected, one-week ZnR strongly decreased fecundity, which was not the case for the restriction of other metals (Fig. 4a). ZnR with the synthetic diet using Nippon gene agarose (less Zn contamination) induced a greater effect, suggesting that egg production is sensitive to the dose of dietary Zn (Fig. 4b). The female flies fed with the ZnR diet for a week had smaller ovaries (Fig. 4c,d). This observation indicated that the decreased fecundity is owing to the defective egg production/maturation, but not to the oviposition behaviour.

**Fig. 4.**
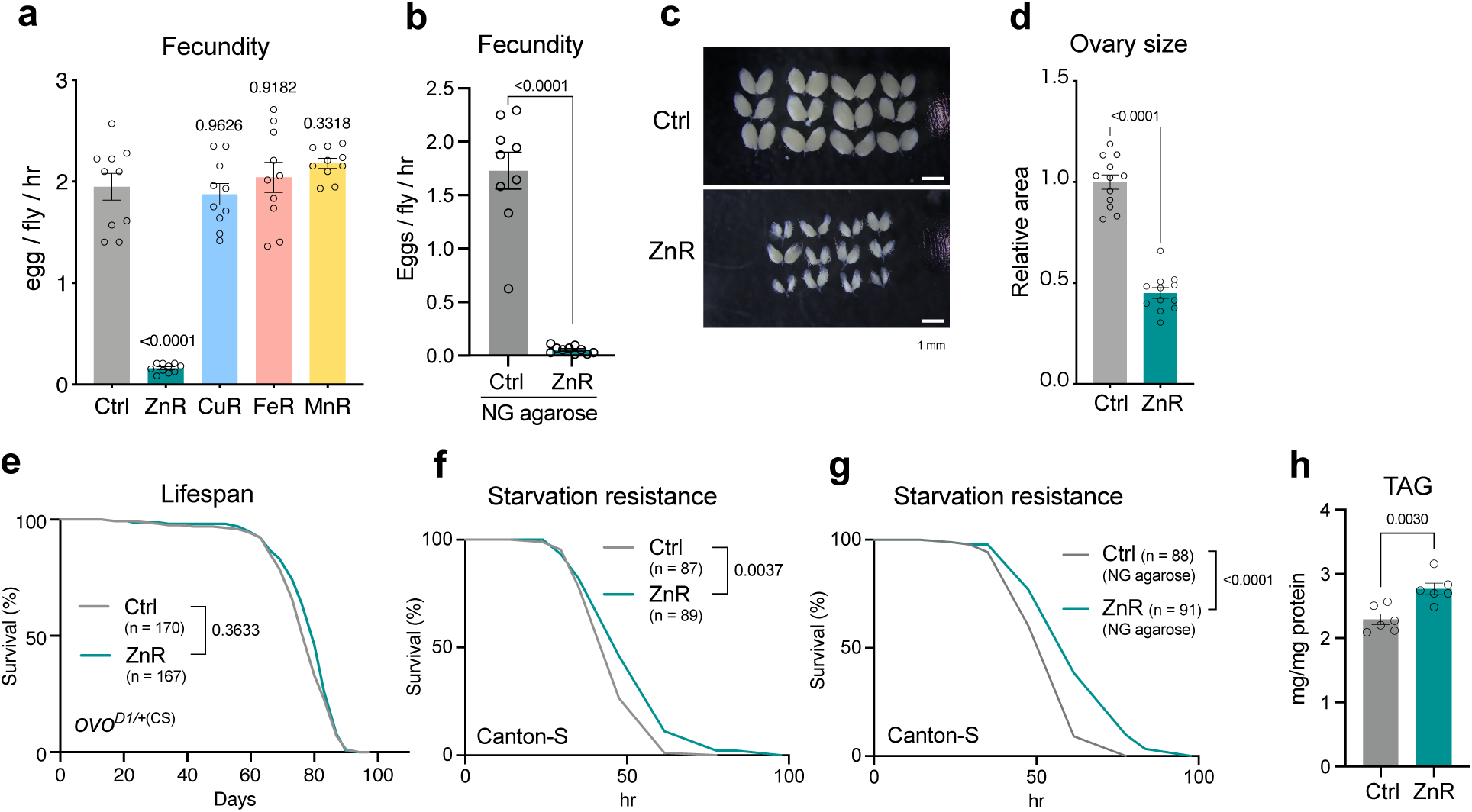
Zinc restriction mimics amino acid restriction phenotypes. **a,** Fecundity of Canton-S flies fed the synthetic diet with or without metal restrictions. n = 10. For the statistics, one-way ANOVA with Dunnett’s multiple comparison test was used. **b,** Fecundity of Canton-S flies fed with or without a zinc-restricted diet using Nippon gene agarose instead of FUJIFILM agar. n = 9 (Ctrl) or 10 (ZnR). For the statistics, a two-tailed Student’s *t* test was used. **c,d,** Representative images (**c**) and the quantification (**d**) of ovaries of Canton-S flies fed with or without a zinc-restricted diet. n = 12. For the statistics, a two-tailed Student’s *t* test was used. Scale bars: 1 mm. **e,** Lifespans of female *ovo*^*D1*/+^ flies fed with or without a zinc-restricted diet. Sample sizes (n) are shown in the figure. For the statistics, a log-rank test was used. **f,** Survivability of female Canton-S flies upon complete starvation after feeding with or without a zinc-restricted diet for one week. Sample sizes (n) are shown in the figure. For the statistics, a log-rank test was used. **g,** Survivability of female Canton-S flies upon complete starvation after feeding with or without a zinc-restricted diet using Nippon gene agarose instead of FUJIFILM agar for one week. Sample sizes (n) are shown in the figure. For the statistics, a log-rank test was used. **h,** The level of triacylglycerol in the female Canton-S flies fed with or without a zinc-restricted diet. For the statistics, a two-tailed Student’s *t* test was used. For all graphs, the mean and SEM are shown. Data points indicate biological replicates. Source data are provided as a Source Data file.

Importantly, lifespan extension by ZnR was not observed in a heterozygous *ovo^D1^* mutant, in which egg production is compromised, suggesting that reduction of fecundity is required for ZnR-longevity (Fig. 4e). We also tested other physiological traits induced by the typical dietary restriction, such as starvation resistance and lipid accumulation. The flies fed with ZnR diet for a week showed increased starvation resistance and lipid storage (Fig. 4f-h).

### Zinc restriction increases feeding preference to yeast

Restriction of dietary yeast increases preference for feeding yeast over sugar^11^. To test whether ZnR changes feeding behaviour, we performed two-choice assay using yeast and sugar. The female flies fed with the ZnR diet for one week showed the preference for yeast (Fig. 5a). Intriguingly, this effect is not due to the increased preference for Zn contained in yeast, as they did not show any augmented preference for Zn-contained sugar over sugar only diet (Fig. 5b). In contrast, they showed increased preference index compared to control when choices were given for the diet contained sucrose but not amino acids versus one contained amino acids but not sucrose, suggesting an increased amino acid preference (Fig. 5c). These data further confirmed that ZnR mimics the dietary yeast restriction phenotype.

**Fig. 5.**
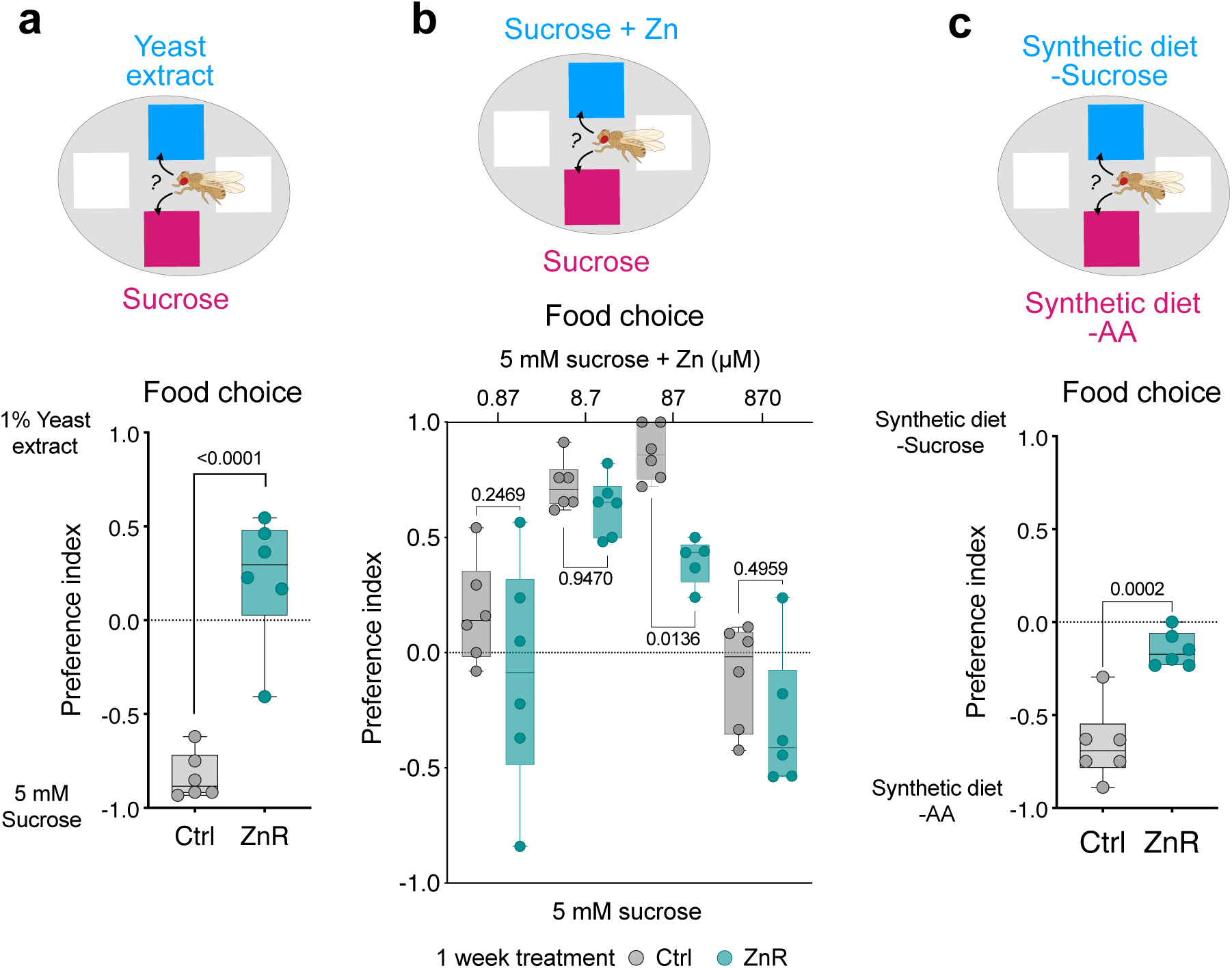
Zinc restriction increases food preference towards yeast rather than sugar. **a,** Food preference towards yeast extract compared to sucrose in female Canton-S flies fed with or without a zinc-restricted diet for 1 week. n = 6. For the statistics, a two-tailed Student’s *t* test was used. b, Food preference towards zinc-supplemented sucrose compared to mere sucrose in female Canton-S flies fed with or without a zinc-restricted diet for 1 week. n = 6. For the statistics, one-way ANOVA with Holm-Šídák’s multiple comparison test was used. **c,** Food preference towards sucrose depleted holidic medium compared to AA depleted holidic medium in female Canton-S flies fed with or without a zinc-restricted diet for 1 week. n = 6. For the statistics, a two-tailed Student’s *t* test was used. For all graphs, the mean and SEM are shown. Data points indicate biological replicates. Source data are provided as a Source Data file.

We noticed that female flies preferred Zn-contained sugar regardless of their dietary environment prior to the choice assay, when the Zn concentration was 87 μM which is equal to the one in the synthetic diet (Fig. 5b). The data suggest that flies can sense Zn and show preference to it. Previous studies revealed that higher concentration of Zn (above 3.3 mM) provokes avoidance as it stimulates some taste neurons such as bitter neurons^12,13^. Thus, we assumed that physiological concentration of Zn (such as in the synthetic diet) can have ability to stimulate sweet or other taste neurons to trigger the attractive behaviour.

## Discussion

In this study, we identified dietary Zn as a regulator of fecundity, starvation resistance, feeding preference, and lifespan. ZnR can phenocopy dietary restriction of protein/amino acids, even under the condition where they have full set of nutrients including twenty amino acids. Thus, Zn is a key micronutrient to determine lifespan-fecundity trade-off, which is consistent with the recent work^14^. Considering the fact that the trade-off is necessary for lifespan extension by ZnR, the decreased fecundity is a first event to trigger lifespan extension. The decreased fecundity in flies with ZnR can be attributed to both the indirect (through IIS) and the direct impact of Zn on oocyte maturation^15^. During oogenesis in *Drosophila*, Zn contents are increased and enriched to form granules in the mature oocyte. Zn is then decreased and distributed uniformly upon the egg activation^15^. The requirement of Zn for oocyte maturation and egg activation is also reported in mammals, including humans^16–18^. The regulation of Zn levels in oocytes, as well as in other tissues, requires Zn transporters, thus manipulation of Zn levels specifically in cells or tissues in the body would be necessary to narrow down the responsible cells for ZnR longevity.

Lifespan extension by the dietary Zn depletion observed in the adult fly was unexpected, given that Zn is an essential nutrient required for biological processes, such as immune, metabolic, or tissue homeostasis. Indeed, in humans, even a mild Zn deficiency (intake of 3.0-5.0 mg/day Zn, instead of 12 mg/day for a control) leads to adverse effects on immunological and biochemical health parameters^19^. Although we did not test all the physiological traits in *Drosophila*, we did not observe striking negative impact on fly health by ZnR, except decreased fecundity. We found that contamination of Zn, such as in agar of the synthetic diet, was sufficient for maintaining a minimum requirement for maintaining health, since animals demand a tiny amount for trace elements, compared to macronutrients such as amino acids. We show that the internal Zn levels during ZnR is approximately 12% of the control. We could not tell whether the zinc level is further decreased during prolonged ZnR or its level is maintained throughout the entire life. However, this level of reduction of internal Zn might not be as severe as it can negatively impact lifespan, at least in *Drosophila*. We assume that adult flies have a way to store Zn obtained during the larval stage and utilise it efficiently throughout the adult life. Of note, the Zinc is reported to be enriched and stored in the Malpighian tubules^20^. There must be mechanisms to augment the absorption of Zn and to block the excretion of it to maintain its level, probably by sensing the decreased level of internal or dietary Zn. It is known that Zn transporters Zip44C.1 and Zip44C.2 in the gut are responsible for Zn absorption, the gene expression of which are regulated by dietary Zn^21^. In the Malpighian tubules, a Zn transporter Zip71B regulates Zn excretion and its expression is also known to be responsive to the dietary Zn levels^22^. Regulatory mechanisms of these Zn transporters by internal Zn levels and the effect of their manipulation on lifespan are yet to be elucidated.

The mechanism of how ZnR extends lifespan is not entirely understood. More than 2800 proteins, reaching 10% of the human genome, are potential Zn proteins^23^. Decrease in cellular Zn levels can therefore influence various processes. One of such mechanisms would be a Zn-gated chloride channel pHCl-2 (also known as Hodor), which regulates IIS through Ilps secretion triggered by TOR signalling in the gut enterocytes^24^. Alternatively, insulin stability and secretion from the mammalian pancreatic β cells are Zn-dependent^25^, although the requirement of Zn for *Drosophila* Ilps might not be conserved. Beyond IIS, there are 302 putative Zinc finger transcription factors in *Drosophila* genome (flybase.org), such as GATA transcription factors which are known to be involved in DR longevity^26^. It is known that age-related activation of innate immunity is one of the hallmarks of ageing, which is related to organismal lifespan^9,27^. Since Zn is tightly linked with immunity^28^, suppression of immune signalling by ZnR would contribute to the lifespan extension.

As far as we know, this study provides the first evidence that limiting dietary Zinc availability by utilising a synthetic diet can extend organismal lifespan. However, it is noteworthy that reducing Zn levels achieved by a zinc-specific chelator TPEN can increase lifespan in *C. elegans*^29^. This manipulation led to the activation of Daf-16, the FoxO ortholog in *C. elegans*, although the lifespan extension was not completely blocked by the loss of *daf16* function alone^29^. It is pivotal to test whether Zinc restriction can activate FoxO to extend lifespan in Drosophila and in mammals.

## Methods

### *Drosophila* stocks and husbandry

Flies were reared on a standard yeast-based diet containing 4.5% cornmeal (NIPPN CORPORATION), 6% brewer’s yeast (ASAHI BREWERIES, HB-P02), 6% glucose (Nihon Shokuhin Kako), and 0.8% agar (Ina Food Industry S-6) with 0.4% propionic acid (Wako 163-04726) and 0.15% butyl p-hydroxybenzoate (Wako 028-03685). For dietary manipulations, we used the exome-matched version of a synthetic diet, or holidic medium, with some modifications^30,31^. Adult flies for experiments were maintained under 25°C, 65% humidity with 12h/12h light/dark cycles. To allow synchronised development and constant density, embryos were collected up to over night using agar plates (2.3% agar, 1% sucrose, and 0.35% acetic acid) with live yeast paste. The equal volume of collected embryos were spread onto plastic bottles. The fly lines used in this study were Canton-S and *ovo^D1^* (Bloomington Drosophila Stock Center (BDSC) 1309).

### Lifespan analysis and starvation resistance assay

Adult flies eclosed within one day were collected and maintained for an additional two days, in order for female flies to mature and mate with males, on the standard yeast-based diet. Flies were then sorted by sex and maintained at a fixed density (30 flies/vial). For lifespan analysis, the number of dead flies was counted every three to four days when flies were transferred to fresh vials. For each lifespan curve, at least six vials were used in parallel to minimise inter-vial variation. For starvation resistance assay, female flies were transferred to vials with 1% agar, and the number of dead flies was counted 2-3 times per day until all flies were dead.

### Inductively coupled plasma-mass spectrometry

Metal contents in flies or in agar (FUJIFILM 010-15815) or agarose (SIGMA A9539-10G, Fast Gene NE-AG01, or Nippon Gene 316-01191) were analysed by ICP-MS (Agilent 7700x). Frozen samples were weighed and placed in a quartz glass test tube. Then 1 mL of 65% HNO_3_ was added and boiled in a water bath at 100°C for eight minutes. After cooling, the volume was fixed at 10 mL with ultrapure water. These solutions were diluted and introduced into an ICP-MS to determine the concentration of metal elements in the eluate.

### Quantification of intestinal stem cell proliferation

Whole guts from female flies were dissected in PBS and fixed in 4% PFA for 1 hr. After washing with PBST (0.1% Triton-X100), the guts were incubated with blocking buffer (PBST (PBS with 0.1% Triton-X100) with 5% normal donkey serum) for 30 minutes. The guts were incubated with anti-Histone H3 (phosphor S28) (1:1000, rat, Abcam, ab10543) antibody diluted in blocking buffer overnight at 4 °C. After washing with PBST, guts were incubated with anti-rat IgG 488 (1:500, Thermo Fisher Scientific, A32766) diluted in blocking buffer for two hours at room temperature. After washing, the guts were mounted using SlowFade Gold (Thermo Fisher Scientific, S36936). Hoechst 33342 (0.4 mM for 1:100, Thermo Fisher Scientific, H3570) was used for nuclear visualisation. The number of phospho-Histone H3 positive cells in the whole gut was counted manually.

### Fecundity analysis

Flies reared on the standard yeast diet until day two post eclosion were maintained for one week in vials with the synthetic diet with 17 females and 17 males. During this period, adult flies were transferred to fresh vials every three days. Flies were then quickly anaesthetised and separated into groups of three males with three females. After 24 hours, the egg number in each vial was counted manually.

### Measurement of triacylglycerol

Four female or male flies were homogenised in PBST (PBS with 5% TritonX-100), then incubated for 5 min at 70°C. Using triglyceride reagent (Wako 632-50991), free glycerol amount derived from triacylglycerol was measured according to the manufacturer’s instruction, based on A_600_. The absorbance was determined by Infinite M Plex (Tecan). Total protein concentration in the same sample was quantified using BCA reagent mix (Wako, 164-25935) for the normalisation.

### Food preference assay

A two-choice test on a Petri dish was modified from the previously described one^32^. Four pieces of square filter paper (2 cm × 2 cm, ADVANTEC No.50) were placed on the bottom of a Petri dish (90 mm diameter). Two pieces in diagonal position were soaked with 150 µl distilled water to humidify the dish. Then, 150 µl of one taste solution (coloured with the red dye, 0.25 mg/mL acid red 52, Wako 018-10012) was applied to one of the remaining pieces of paper, and the other taste solution (coloured with the blue dye, 0.125 mg/mL brilliant blue, Wako 027-12842) was applied to the other. For the two-choice assay using yeast and sugar, 1% yeast extract and 5 mM sucrose were coloured blue and red, respectively. To assess Zn preference, 5 mM sucrose with 0.87 μM (1/100×), 8.7 μM (1/10×), 87 μM (1×), or 870 μM (10×) ZnSO_4_ solution and 5 mM sucrose were coloured blue and red, respectively. To examine AA preference, agar, cholesterol, propionic acid, and nipagin were excluded from the holidic medium solution. Holidic medium solution from which 20 AAs were omitted and holidic medium solution from which sucrose was omitted were coloured red and blue, respectively.

Adult female flies were allocated to vials containing a synthetic diet at day two post eclosion. Six vials, each containing 30 individuals, were prepared for each condition. After one week, flies were transferred to vials containing 1% agar for starvation for 28 hours. The starved flies were then quickly anesthetised by exposure to a temperature of 4°C for 10 minutes and subsequently transferred to the assay plate. The plate was covered by aluminium foil and positioned in the incubator at 25°C with a humidity level of 60% for two hours. Subsequently, the plate was retrieved and subjected to freezing at −30°C for at least two hours. The coloration of the abdomen was manually categorised into blue, purple, and red. The preference index (PI) towards yeast was calculated as follows: PI[blue]={N(blue)-N(red)}/{N(blue)+N(purple)+N(red)}. Vials with more than 30% colorless flies were excluded from the analysis.

### Quantification and statistical analysis

Statistical analysis was performed using Graphpad Prism 9 and 10. A two-tailed Student’s *t*-test was used to test between samples. One-way ANOVA with Holm-Šídák’s multiple comparison test was used to test among groups. One-way ANOVA with Dunnett’s multiple comparison test was used to test between control and multiple samples. Bar graphs were drawn as mean and SEM with all the data point shown by dots to allow readers to see the number of samples and each raw data. OASIS 2 was used to perform log-rank test for lifespan analysis^33^.

## Acknowledgements

We would like to acknowledge Taiho Kambe, Yoriko Akuzawa-Tokita, and all the Obata lab members for the technical assistance and valuable suggestions. We thank National Institute of Genetics, Kyoto Drosophila Stock Center, Vienna Drosophila Resource Center, and Bloomington Drosophila Stock Center for fly stocks. This work was supported by AMED-PRIME to F.O. (JP20gm6310011), and partly by AMED-Project for Elucidating and Controlling Mechanisms of Aging and Longevity to M.M. This work was also supported by grants from the Japan Society for the Promotion of Science to F.O. (22H02769).

## Author Contributions

F.O. and H.K. conceived the project. H.K., H.A., R.O., performed most of the experiments and analysed the data. S.K. quantified ISC proliferation and performed food choice assay. C.S. established the method for food choice assay. H.K. and F.O. wrote the initial manuscript. M.M. and F.O. supervised the study. All authors edited and approved the final manuscript.

## Competing interests

The authors declare no competing interests.

## Materials & Correspondence

All the materials generated in this study are available upon request to F.O.

## Notes

### Competing Interest Statement

The authors have declared no competing interest.

### Summary of Updates

Discussion updated and a citation added

